# The bacterial pathogen causing cucurbit yellow vine disease associates with the specialized extrafascicular phloem unique to cucurbits

**DOI:** 10.64898/2026.09.24.754277

**Authors:** Kephas Mphande, Breah LaSarre, Mark L. Gleason, Gwyn A. Beattie

## Abstract

Cucurbit yellow vine disease (CYVD) is a phloem disease caused by a group of *Serratia ureilytica* bacteria within the *S. marcescens* complex. The CYVD pathogen is distinct from other phloem pathogens in two fundamental ways: i) unlike other bacterial phloem pathogens, the CYVD pathogen readily grows on standard laboratory media; and ii) whereas most bacterial phloem diseases are transmitted solely by hemipteran insects, CYVD can be transmitted by both hemipteran (squash bug) and non-hemipteran (cucumber beetle) vectors. The phloem of cucurbits is also distinct, with cucurbits having a unique secondary phloem system, the extrafascicular phloem (EFP), for which a role in CYVD has not been investigated. The distinctive features of this pathosystem prompted a re-evaluation of pathogen localization *in planta* using a green fluorescent protein-tagged CYVD strain during infection of squash (*Cucurbita pepo*). In epifluorescence and confocal microscopy images of transverse and longitudinal sections of stem and petiole tissues, the fluorescence patterns of the CYVD pathogen were consistent with pathogen colonization of all four known EFP tissues, namely the peripheral, entocyclic, ectocyclic, and commissural sieve tubes. The CYVD pathogen may enter the EFP during herbivory by pathogen-infected vectors that cause mechanical damage, as the EFP exhibits much slower blockage following injury than the conventional fascicular (bundle) phloem. Moreover, although EFP has been proposed to have an antimicrobial function, EFP exudate collected from the stem and petioles of *C. pepo* supported rapid pathogen growth. These results provide evidence for a unique tissue tropism of the newly emerged plant pathogen *S. ureilytica*.

**SIGNIFICANCE STATEMENT:** Bacterial phloem pathogens generally colonize the conventional fascicular (bundle) phloem of their host plant. We demonstrate that the bacterial pathogen causing cucurbit yellow vine disease is atypical in that it colonizes the extrafascicular phloem, a unique secondary phloem system of cucurbits that is thought to function beyond photosynthate transport. These findings reshape our understanding of phloem pathogenicity and indicate that the extrafascicular phloem is more physiologically accessible and biologically consequential than previously recognized.

## INTRODUCTION

Cucurbit yellow vine disease (CYVD) is an emerging phloem disease affecting a variety of cucurbit crops, including pumpkins and squash (*Cucurbita spp.*), watermelon (*Citrullus lanatus*), and cantaloupe and muskmelon (*Cucumis melo*) (Bruton et al. 1998; Bruton et al. 2003). CYVD is characterized by phloem tissue discoloration, particularly in the crown region of the stem, and also by leaf yellowing, vine decline, and stunting, with onset often during the fruit development period (Bruton et al. 1995; Bruton et al. 1998). Reports of CYVD are restricted to the United States, with the first report in Texas and Oklahoma in 1988 (Bruton et al. 1995) and subsequent reports indicating a range extending from Texas to Massachusetts (Bost et al. 1999; Wick et al. 2001; Rascoe et al. 2003; Sikora et al. 2012; Besler and Little, 2015; Rodriguez et al. 2023; Mphande et al. 2024).

CYVD was originally identified as a phloem disease, even before identification of the causal agent, based on safranin staining of the vascular tissue and observation of “bacteria-like-organisms” by transmission electron microscopy in the sieve elements of symptomatic cucurbit plants (Bruton et al. 1998). Bacterial phloem pathogens are rare and include the cell wall-less *Candidatus* Phytoplasma spp., multiple *Ca.* Liberibacter spp., *Ca.* Arsenophonus phytopathogenicus*, Ca.* Phlomobacter fragariae (corrected to *Ca.* Phloeobacter fragariae (Oren 2017) and several *Spiroplasma* spp. These pathogens live intracellularly within the sieve elements, binding to the plasma membrane and endoplasmic reticulum of their host cells (Nome et al. 2009; Hartung et al. 2010; Buxa et al. 2015; van Bel and Musetti 2019; Achor et al. 2020; Dittmer et al. 2021). Nearly all phloem pathogens have candidate genus status because they are recalcitrant to cultivation on laboratory media.

CYVD is caused by a subset of strains of *Serratia ureilytica*, a species within the *S. marcescens* complex (Avila et al. 1998; Rascoe et al. 2003; Mphande et al. 2025b). Studies over the past several decades have shown that the CYVD pathogen differs from other bacterial phloem pathogens in two fundamental ways. First, CYVD-inducing *S. ureilytica* (hereafter referred to as CYVD strains) grow robustly on standard laboratory media, in contrast to the unculturable nature of all other phloem pathogens except *Spiroplasma* spp., which grow on highly specialized media (Lee and Davis 1989). Second, the CYVD pathogen is the only bacterial phytopathogen that can be transmitted by both a hemipteran, namely squash bugs (*Anasa tristis*) (Bruton et al. 2003; Pair et al. 2004), and non-hemipterans, namely the cucumber beetle species *Acalymma vittatum* and *Diabrotica undecimpunctata howardi* (Mphande et al. 2025b). These insects cause extensive mechanical damage to plants during feeding. Squash bugs damage plant tissues using their large stylets and probing feeding style (Mitchell 2004), and cucumber beetles damage plants through chewing herbivory. In contrast, other phloem pathogens are transmitted solely by hemipterans that cause minimal plant damage by navigating their thin stylets along an intercellular pathway to individual sieve elements (Sharma et al. 2014). The subsequent piercing-sucking action can move a vector-associated pathogen into the sieve elements against the strong outward force generated by phloem sap. Collectively, these distinctions suggest that interactions between CYVD strains and the phloem may differ from those of other phloem pathogens.

The phloem structure of cucurbits is atypical as well. Cucurbits have bicollateral vascular bundles, with conventional photoassimilate-transporting phloem, called fascicular (bundle) phloem, located both internal and external to the xylem (Figure 1). The Cucurbitaceae family is unique in also having a second phloem system, called the extrafascicular phloem (EFP) (Crafts 1932; Turgeon 2016). In *Cucurbita pepo,* the EFP comprises three components that are longitudinal in the stems and petioles: the peripheral sieve tubes, which lie in an arc at the periphery of the internal and external fascicular phloem, and the entocyclic and ectocyclic sieve tubes, which lie inside and outside of the sclerenchyma ring, respectively. The EFP also includes a fourth component, the commissural sieve tubes, which run transversely (horizontally) through the cortex of the stems and petioles.

**Figure 1.**
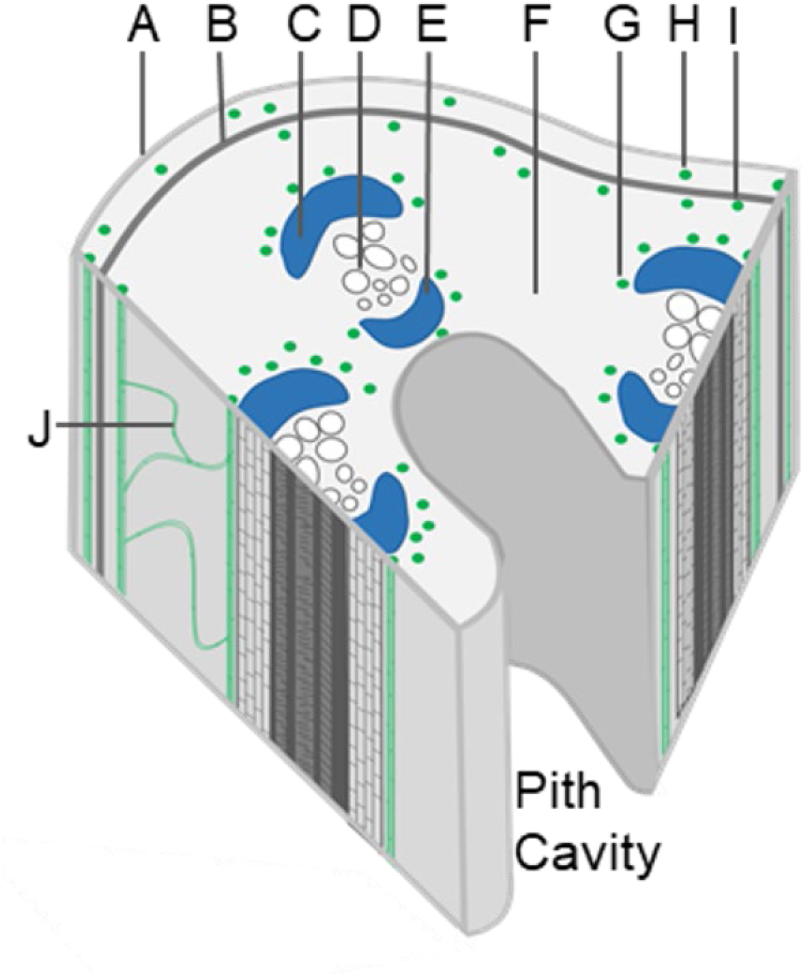
**Annotated schematic of the major vascular and nonvascular tissues in stem section of *Cucurbita pepo.*** The fascicular (bundle) phloem is colored blue, and extrafascicular phloem (EFP) types are colored green. A, epidermis; B, sclerenchyma ring; C, external fascicular phloem; D, xylem; E, internal fascicular phloem; F, cortical parenchyma; G, peripheral EFP; H, ectocyclic EFP; I, entocyclic EFP; J, commissural EFP.

In this report, we provide evidence that the CYVD pathogen localizes to the four component tissues of the EFP. While the primary function of the fascicular phloem is the transport of photoassimilates (Zhang et al. 2010; Kanvil et al. 2016; Schnieder et al. 2022), current evidence suggests that the EFP functions in the transport of other nutrients (Tolstikov and Fiehn 2002; Chen et al. 2004; Zhang et al. 2010) and/or in signaling and defense (Zhang et al., 2010; Gaupels et al. 2012; Kanvil et al. 2016). Our results highlight that the EFP is not only physiologically accessible to microbes via insect herbivory, but also biologically consequential in its conduciveness to colonization by an insect-vectored bacterial pathogen.

## RESULTS

Considering the distinct CYVD strain biology and cucurbit phloem architecture relative to other phloem pathosystems, we re-evaluated the tissue tropism of the CYVD pathogen *in planta* using a green fluorescent protein (GFP)-expressing derivative of CYVD strain Z07, hereafter referred to as Z07(pGFP). We used summer squash as a model cucurbit because, along with pumpkin, squash exhibits particularly high disease incidence and crop losses from CYVD (Cartright and Bruton 1993; Bruton et al. 1998; Bruton et al. 2003). Z07(pGFP) infection caused disease symptoms comparable to those caused by wild-type Z07 (Figure S1).

### *S. ureilytica* localizes to regions peripheral to the fascicular phloem

Pathogen localization was assessed 14 d after inoculation by epifluorescence microscopy and confocal microscopy of hand-sectioned stems and petioles. To demarcate regions of phloem tissue, we performed live-plant phloem labeling with carboxyfluorescein diacetate, a membrane-permeable precursor that is phloem-mobile and hydrolyzed intracellularly into the fluorescent carboxyfluorescein (CF) tracer. When introduced via foliar abrasion, CF most intensely labels the conventional fascicular phloem of adjacent petioles but can also label the EFP after longer staining periods (Zhang et al. 2010).

The localization patterns of the CF tracer and Z07(pGFP) differed markedly. In petiole transverse sections, CF exhibited the characteristic pattern for bicollateral fascicular phloem (Figure 2a, red arrows). Specifically, the CF tracer was concentrated in crescent-shaped regions flanking the inner and outer margins of the xylem. In transmitted-light images, these regions typically had a hazy appearance and were darker than the adjacent parenchyma but lighter than the xylem (Figure 2a, bright field). The CF tracer was occasionally visible extending outward from the vascular bundles in the commissural sieve tubes of the EFP (Figure 2a, blue arrowheads), as previously seen in cucumber and pumpkin (Zhang et al. 2010; Sui et al. 2021). In contrast, Z07(pGFP) localized in scattered foci along arcs that were near but peripheral to the internal and external fascicular phloem (Figure 2b,c). Mock-infected controls exhibited negligible fluorescence (Figure 2d; Figure 3), and the lifetime of the fluorescent signal (τ) matched that of Z07(pGFP) cells grown in pure culture (τ = 1.7 ns), validating that the fluorescence originated from Z07(pGFP) cells. The arcs of Z07(pGFP) foci varied in shape and density among vascular bundles but were typically broad, with many foci near the fascicular phloem. Similar localization was observed in petioles and areas of the stem both above and below the inoculation site (Figure 3). The presence of Z07(pGFP) foci in arcs around the periphery of the fascicular phloem was also observed in a recent paper (Rodriguez-Herrera et al. 2026) and matches the distribution pattern of peripheral EFP sieve tubes as identified using immunostaining of EFP-localized proteins (Smith et al. 1987; Bostwick et al. 1992; Schnieder et al. 2022). These results are consistent with localization of this pathogen in or near the peripheral sieve tubes of the EFP.

**Figure 2.**
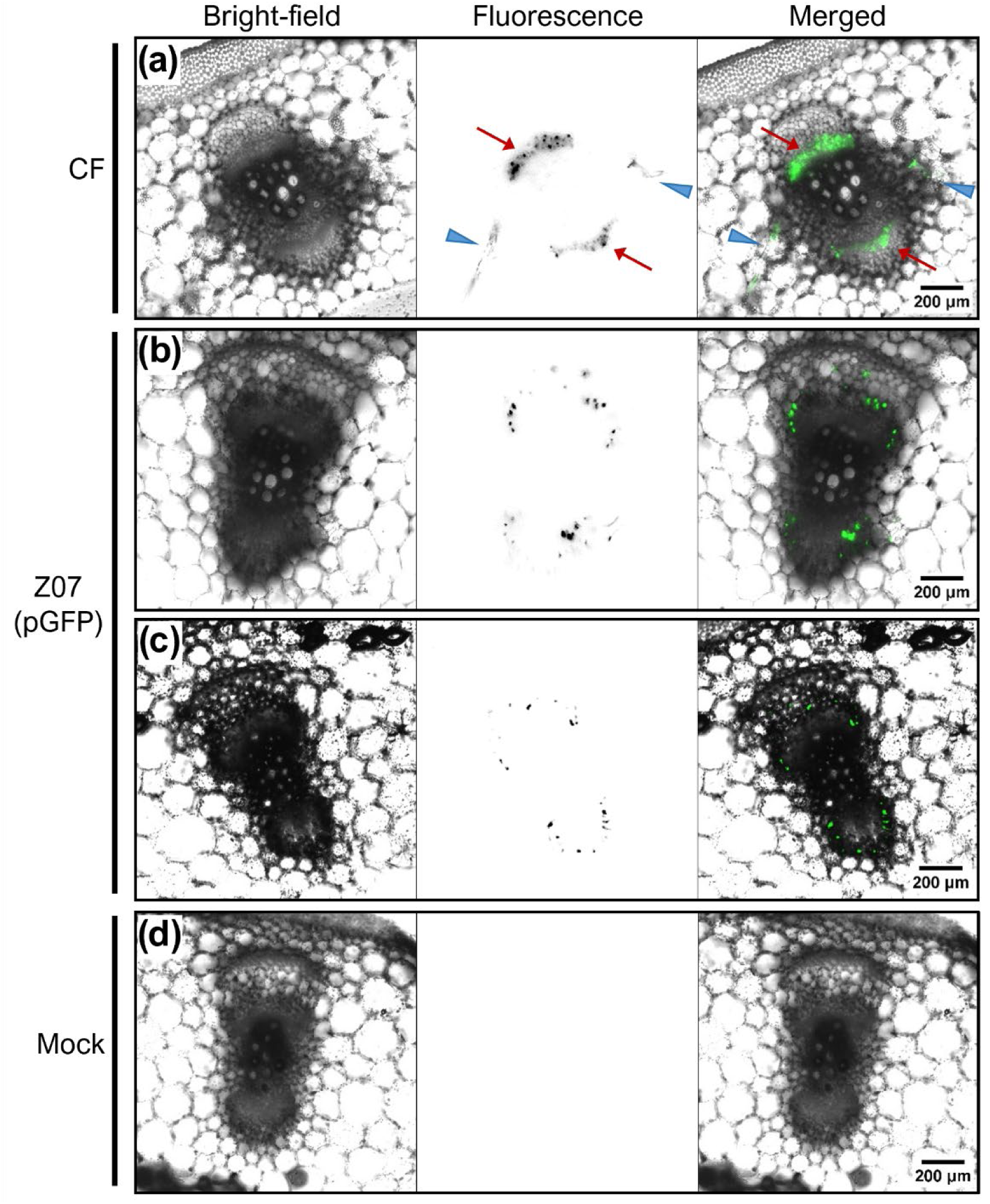
Z07(pGFP) localizes around the perimeter of the vascular bundles in *C. pepo* petioles. (a) Epifluorescence microscopy images of a petiole stained with the fluorescent phloem-mobile tracer carboxyfluorescein (CF). Arrows indicate CF fluorescence in fascicular phloem (red arrows) and commissural extrafascicular phloem (EFP) (blue arrowheads). (b, c) Petiole cross-sections from Z07(GFP)-infected plants imaged by (b) epifluorescence microscopy and (c) confocal microscopy. (d) Epifluorescence microscopy images of a petiole from a mock-inoculated control plant. Identical brightness and contrast adjustments were applied to the fluorescence images in (b) and (d). (a–d) Fluorescence is shown as an inverted grayscale image in the fluorescence-only panels and in green in the merged panels.

**Figure 3.**
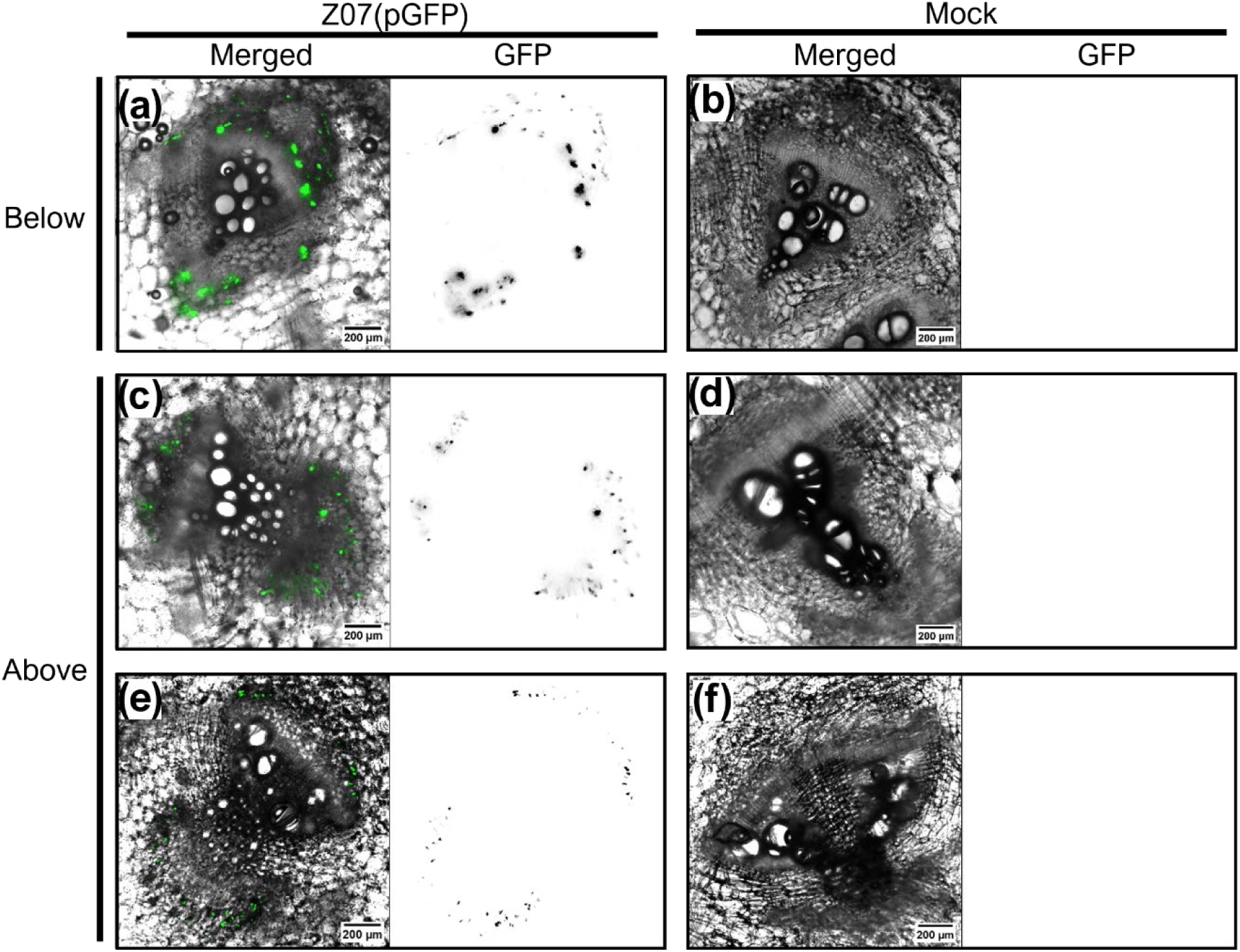
Z07(pGFP) localizes around the perimeter of the vascular bundles in *C. pepo* stems. (a–d) Merged bright-field/GFP epifluorescence microscopy images and corresponding GFP channel alone of stem cross-sections from below the inoculation site (a, b) or above the inoculation site (c, d) of an Z07(GFP)-infected plant (a, c) or mock-inoculated plant (b, d). (e, f) Merged transmitted light/GFP confocal microscopy image and corresponding GFP channel alone of a stem cross-section from above the inoculation site of an Z07(GFP)-infected plant (e) or mock-inoculated plant (f). For all panels, the fluorescence channel alone is shown as an inverted grayscale image. Identical brightness and contrast adjustments were applied to the fluorescence images in each pair of Z07(pGFP)-infected and mock-inoculated cross-sections.

**Transverse sections highlight localization of *S. ureilytica* at sites of known EFP tissues** To investigate the extent of Z07(pGFP) co-localization with EFP, we also examined tissue regions known to support other types of EFP. We consistently observed Z07(pGFP) foci in specific regions away from the vascular bundles (Figure 4). These foci were often sufficiently distant to be excluded from images centered on vascular bundles; moreover, when visible in an image field with Z07(pGFP) foci bordering vascular bundles, the fluorescence intensity of the more distant foci was often lower, making them easy to overlook despite being readily distinguishable from the background. In transverse stem sections, Z07(pGFP) foci were commonly visible immediately internal and external to the subepidermal sclerenchyma ring (Figure 4a,c), which aligns with the known locations of the entocyclic and ectocyclic EFP sieve tubes, respectively (Smith et al. 1987; Kempers et al. 1993; Schnieder et al. 2022). Petioles lack a continuous subepidermal sclerenchyma ring, but clear Z07(pGFP) foci were visible in the subepidermal regions at the interface between the outer ring of photosynthetic cortical parenchyma and either the collenchyma or the epidermis in regions lacking collenchyma (Figure 4b,d). This localization indicates the pathogen associates with ectocyclic or endocyclic EFP in petioles, although the distinction between these two EFP types in petioles remains unresolved (Figure S1). Although captured less frequently, we also observed Z07(pGFP) fluorescence in presumptive commissural sieve tubes (Figure 4c,d).

**Figure 4.**
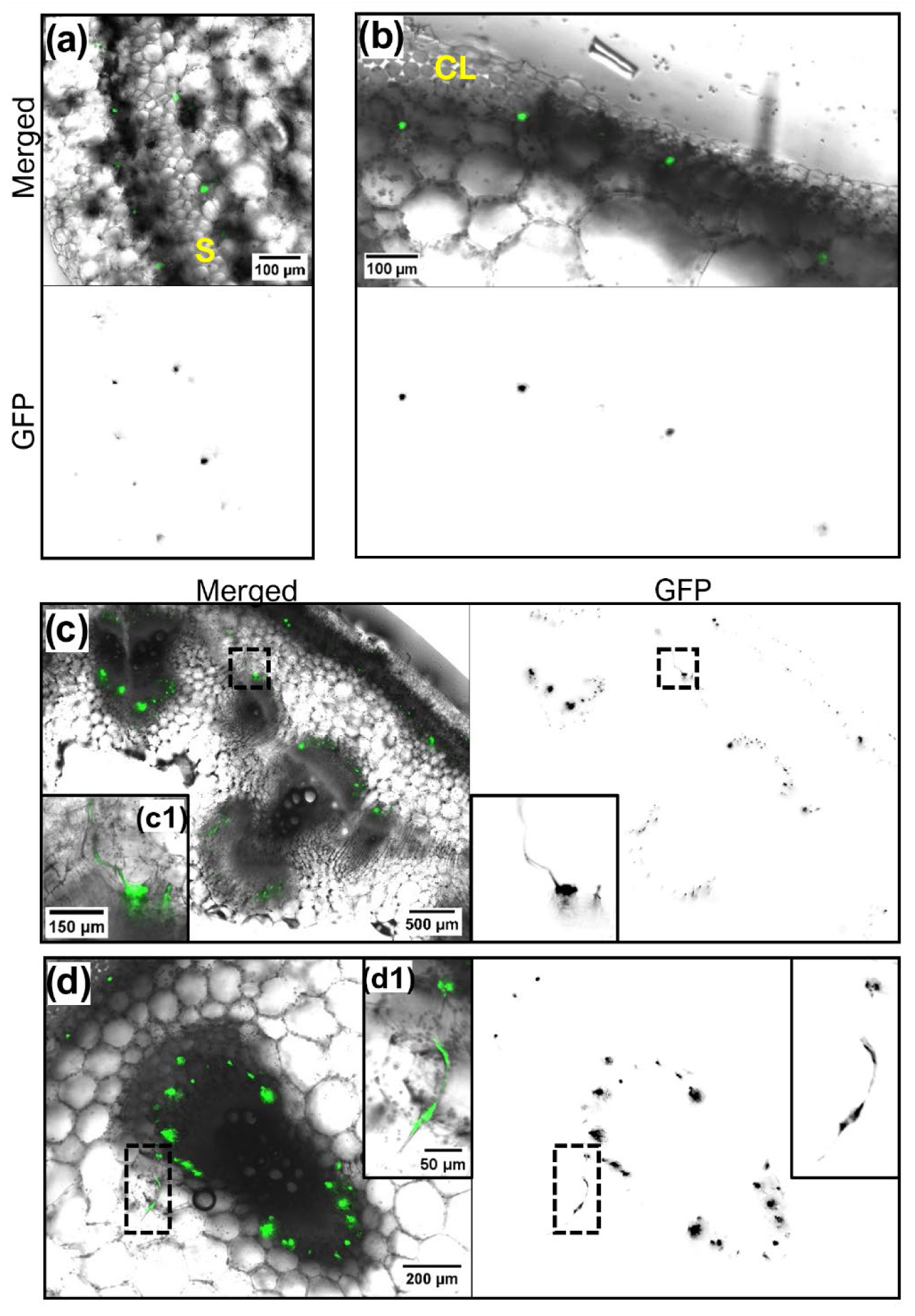
Z07(pGFP) localizes to regions of all four types of EFP in *C. pepo*. Merged bright-field/GFP epifluorescence microscopy images and corresponding GFP channel alone for (a, c) stem and (b, d) petiole cross-sections from Z07(pGFP)-infected squash plants. The GFP channel alone is shown as an inverted grayscale image in all panels. (a) Stem region encompassing the sclerenchyma ring (labeled as S). (b) Petiole region containing part of a collenchyma patch (labeled as CL) and adjacent epidermal region lacking collenchyma. (c) Stem region spanning multiple vascular bundles. (d) Petiole region containing a vascular bundle and collenchyma. (c1, d1) Enlarged dashed-box regions from (c) and (d), respectively, showing lateral strands of Z07(pGFP) corresponding to presumptive commissural EFP.

### Longitudinal sections provide evidence of *S. ureilytica* in EFP sieve tubes

We also analyzed pathogen localization patterns in longitudinal stem sections, imaging sections only if they contained unambiguous xylem that was recognizable by large vessel lumens or annular or spiral secondary wall thickenings, which served as landmarks for tissue identification. Consistent with fluorescence patterns in transverse stem sections, longitudinal stem sections showed that Z07(pGFP) localized to three discrete regions between the pith cavity and epidermis: i) the pith-facing margin of the internal phloem; ii) the epidermis-facing margin of the external phloem; and iii) a region straddling the sclerenchyma ring (Figure 5a). Notably, the GFP fluorescence in longitudinal sections occurred in elongated strands that ran parallel to the stem axis (Figure 5a,d; Figure 6a). Z-stack visualization revealed that strands appearing as short and discontinuous in single-plane images were often segments of longer continuous strands that traversed multiple focal planes (Figure 6c). The sieve elements of EFP are known to be smaller in diameter (≤ 20 μm) than those of the fascicular phloem (typically 50 μm) (Crafts 1932; Smith et al. 1987; Knoblauch and Oparka 2012). Here, the strands were typically 5–15 μm wide (Figure 5b,c,e), consistent with EFP sieve tubes, and spanned distances substantially greater than the length of individual parenchyma cells (Figure 5e; Figure 6b,c). Within some strands, the GFP signal appeared as discrete puncta consistent in size with individual *S. ureilytica* cells (Figure 5b,c). The fluorescent strands were occasionally interrupted by nonfluorescent transverse bands (Figure 7), suggestive of sieve plates. The distribution of the Z07(pGFP) strands strongly aligns with the EFP as mapped in previous studies, including in longitudinal sections of *Cucumis sativus* using immunostaining of the EFP-localized PP1 protein (Schnieder et al. 2022). Thus, as in transverse sections, Z07(pGFP) localization in longitudinal stem sections supported the association of the CYVD pathogen with the EFP rather than the fascicular phloem, and, moreover, indicated its presence within EFP sieve tubes.

**Figure 5.**
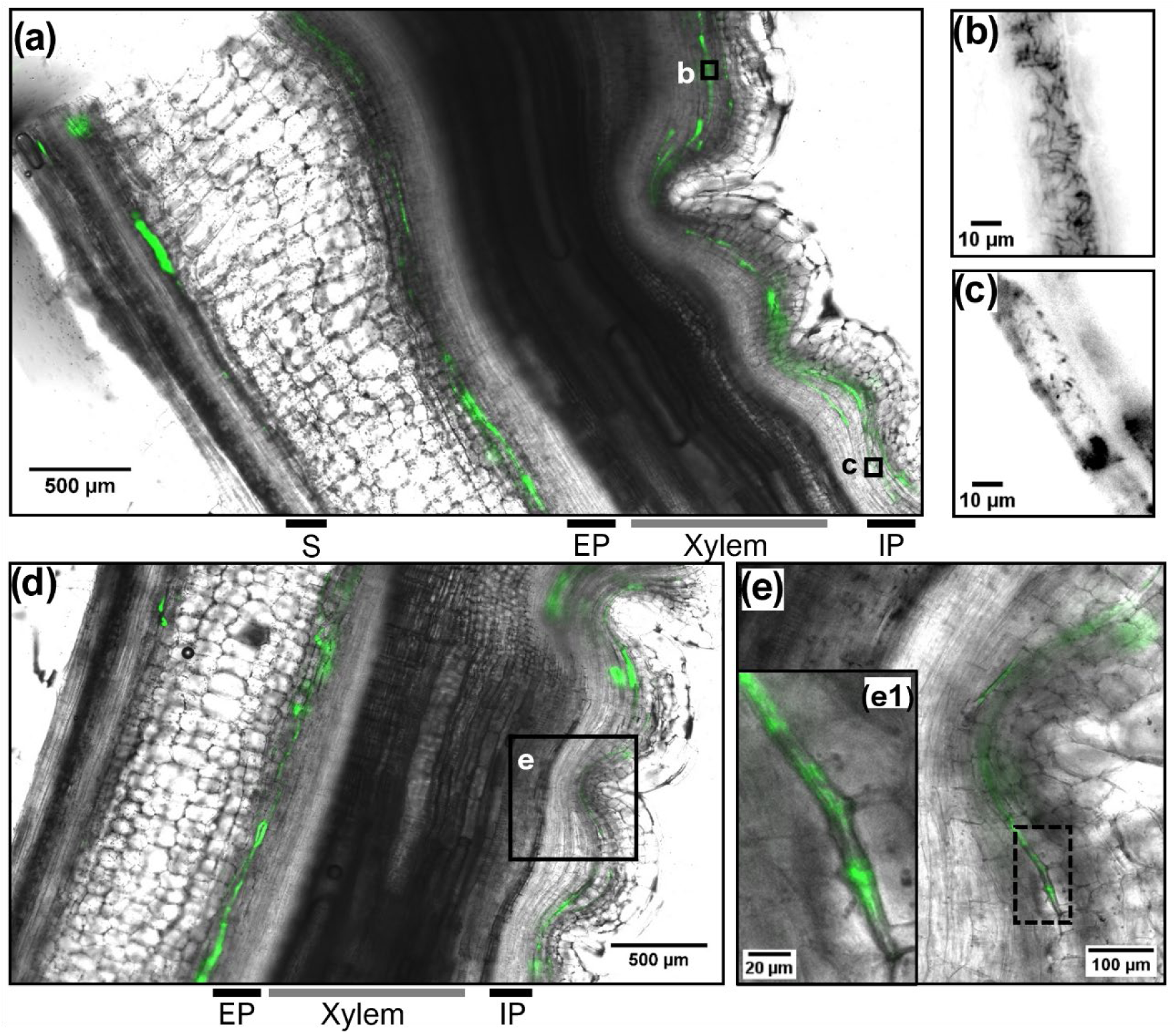
Z07(pGFP) forms longitudinally oriented strands in the *C. pepo* stem. (a) Merged bright-field/GFP epifluorescence microscopy image of a longitudinal stem section. Regions of the sclerenchyma ring (S), external fascicular phloem (EP), xylem, and internal fascicular phloem (IP) are indicated below the figure. (b, c) Higher-magnification GFP epifluorescence images of the boxed regions as labeled in (a), shown as inverted grayscale images. (d) Merged bright-field/GFP epifluorescence microscopy image of a longitudinal stem section. Regions of IP, xylem, and EP are indicated below the figure. (e) Higher-magnification merged image of the boxed region as labeled in (d). (e1) Enlarged dashed-box region from (e) showing a fluorescent strand approx. 10 µL wide.

**Figure 6.**
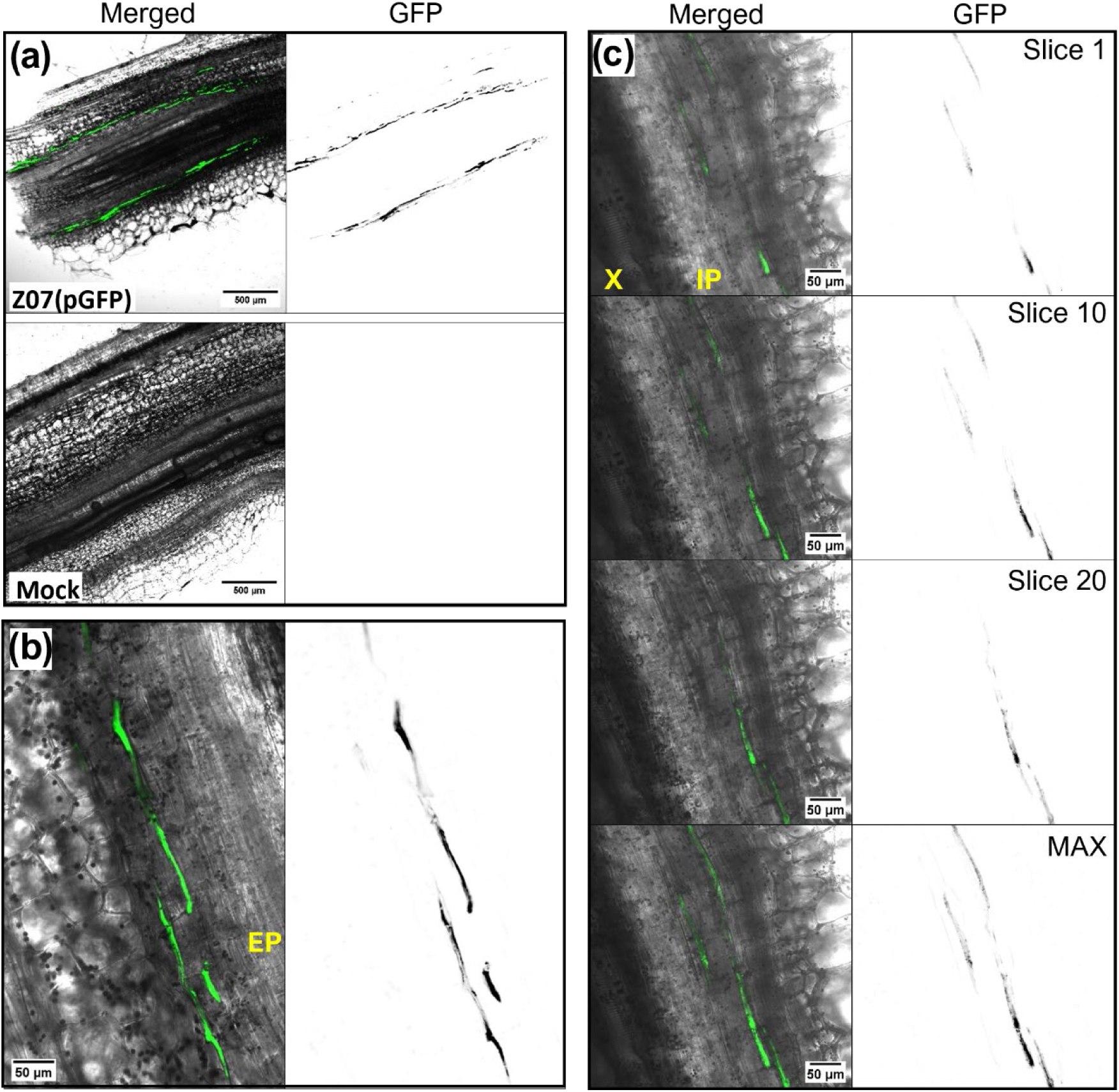
Longitudinal Z07(pGFP) strands span distances greater than the length of individual parenchyma cells. Merged transmitted light/GFP confocal microscopy images and corresponding GFP channel alone of longitudinal stem sections. For all panels, the GFP channel alone is shown as an inverted grayscale image. (a) Images from a Z07(pGFP)-infected plant (top) and a mock-inoculated control plant (bottom). Identical brightness and contrast adjustments were applied to the fluorescence images. (b) Stem region of a Z07(pGFP)-infected plant encompassing the external fascicular phloem (EP) and parenchyma cells. (c) Images from a 20-slice Z-stack (0.91 µm optical section spacing; total depth 17.28 µm) of a Z07(pGFP)-infected stem section. The top three sets of images show individual optical sections (slices 1, 10, and 20). The bottom set of images shows a maximum-intensity projection of the GFP channel from all 20 slices and the corresponding merged image with transmitted light from slice 10. X, xylem. IP, internal fascicular phloem.

**Figure 7.**
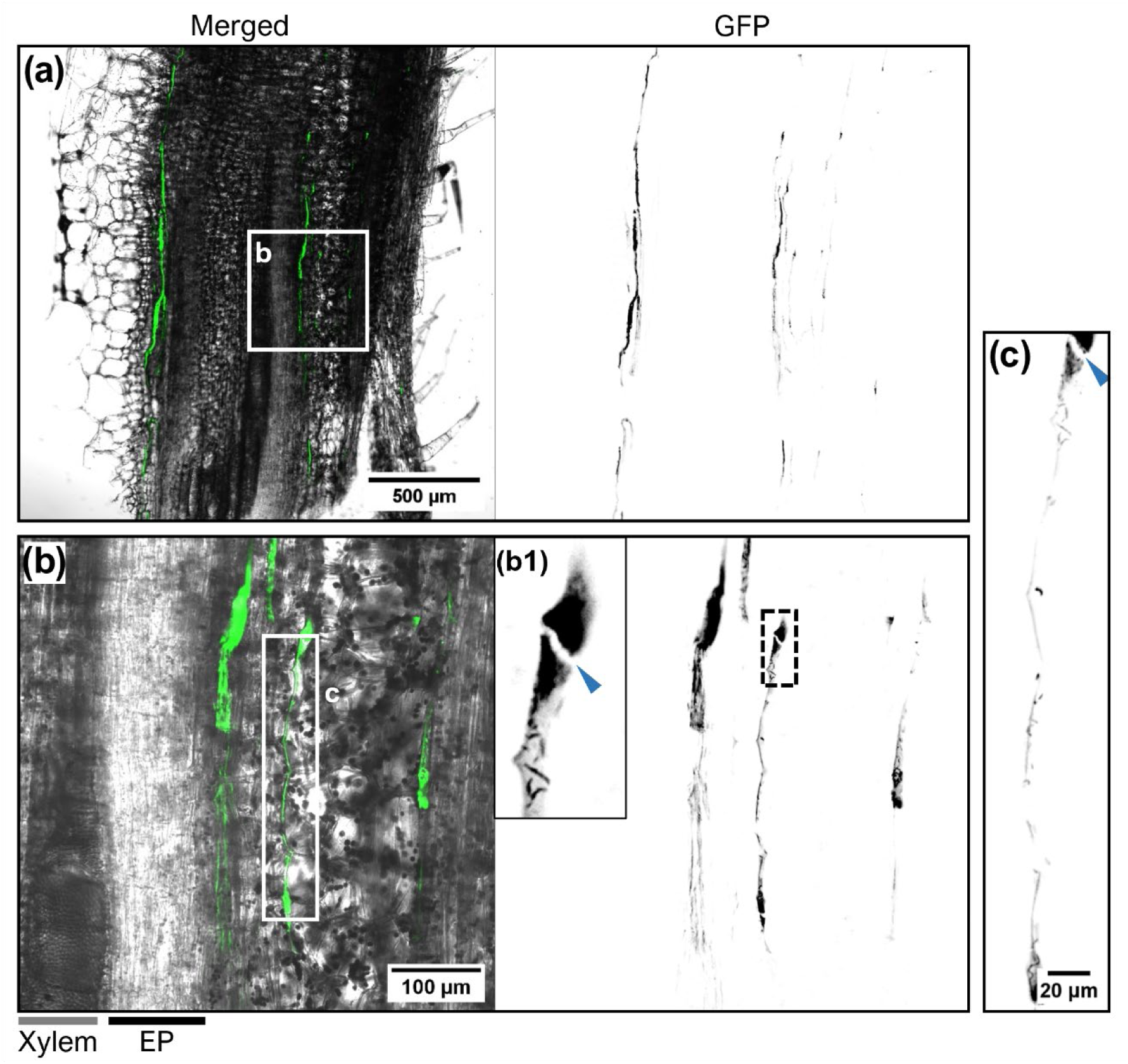
Z07(pGFP) strand fluorescence is occasionally disrupted by a nonfluorescent transverse band, suggestive of a sieve plate. (a) Merged transmitted light/GFP confocal microscopy image and corresponding GFP channel alone of a longitudinal stem section. (b) Higher-magnification merged and GFP images of the boxed region as labeled in (a). Regions of xylem and external phloem (EP) are indicated below the figure. (b1) Enlarged dashed-box region from (b), with arrow indicating the putative sieve plate. (c) Maximum-intensity projection of GFP fluorescence from a 13-slice Z-stack (3.25 µm optical section spacing; total depth 39.1 µm) of the boxed region as labeled in (b), with arrow indicating the putative sieve plate. For all panels, the GFP channel alone is shown as an inverted grayscale image.

### EFP exudate supports growth of the *S. ureilytica* CYVD pathogen

The EFP has been proposed to function in defense against insects and pathogens based on EFP exudate containing secondary metabolites and putative defense proteins (Tolstikov and Fiehn 2002; Turgeon and Oparka 2010; Gaupels et al. 2012; Gaupels and Ghirardo 2013). Moreover, a role in antiherbivory was demonstrated by EFP exudate-induced feeding aversion and reduced growth in aphids (Kanvil et al. 2016); a similar role against pathogens has not been addressed.

Here, we evaluated how EFP exudate from the cut stems and petioles of a healthy squash plant affected the growth of the CYVD pathogen. Cucurbit stem exudates originate primarily from the EFP and not the fascicular phloem because the fascicular phloem undergoes rapid injury-induced occlusion whereas the EFP does not (Zhang et al. 2012; Lopez-Cobollo et al. 2016; Schnieder et al. 2022). The extent to which the EFP exudate is diluted by xylem sap varies with the distance of the cut from the soil (Schnieder et al. 2022) and the plant species and growth conditions (Zhang et al. 2012); therefore, EFP exudates were collected under conditions expected to minimize xylem sap contamination. When cells of the CYVD pathogen Z07 or a mesophyll leaf pathogen, *Pseudomonas syringae* pv. syringae, were introduced into exudates collected from the cut stems and petioles of squash, cells recovered from the exudate after 24 h exhibited altered colony phenotypes, indicating a bacterial response to the exudate (Figure S2b). However, both pathogens grew to high densities in the liquid exudate (Figure S2a). Additionally, one class of secondary compounds, cucurbitacins, which are involved in antiherbivory (Lang et al. 2013) and toxicity to insects (Ponsakar et al. 2020), did not show antimicrobial activity against either of these pathogens when present at a concentration known to inhibit *Phytophthora cactorum* (Nes and Patterson 1981) (Figure S2c). Although we cannot exclude the possibility that antimicrobial compounds in the EFP exudate were unstable, diluted by xylem sap or apoplastic fluids, or required pathogen infection for induction, our collective data do not support an antimicrobial role for the EFP.

## DISCUSSION

Our results demonstrate that *S. ureilytica* CYVD strains are unlike all known bacterial phloem pathogens in that they do not colonize the sieve tubes of fascicular phloem. Instead, the CYVD pathogen associates with the cucurbit-specific EFP at sites both bordering and distant from the vascular bundles. The bundle-associated sites of pathogen localization described herein are consistent with independent observations reported recently (Rodriguez-Herrera et al. 2026), although that study restricted the imaging to vascular bundles, which would have prevented detection of the pathogen in distal tissues. The strands of fluorescence in longitudinal sections of cucurbit stems strongly indicate localization within the EFP sieve tubes, which could serve as long-distance transport channels for systemic movement of the pathogen (Rodriguez-Herrera et al. 2026). However, we cannot discount that the pathogen may also colonize the intercellular spaces, or apoplast, adjacent to the EFP, as suggested by the shape of some fluorescent foci in transverse sections (Rodriguez-Herrera et al. 2026; Figure S3). The biological basis for pathogen localization to EFP regions is intriguing and warrants further investigation.

To date, all known phloem pathogens colonize the conventional fascicular (bundle) phloem of their host plants. Our findings thus broaden our understanding of phloem pathogenicity to include colonization of these functionally and physiologically distinct EFP tissues. We predict that the fascicular and extrafascicular phloem offer distinct chemical environments for a pathogen, as supported by differences in the metabolome and proteome of the EFP versus the fascicular phloem (Gaupels et al. 2012; Lopez-Cobollo et al. 2016). These differences suggest limited mixing of the two phloem systems. That said, the EFP is interconnected with the fascicular phloem via the commissural sieve tubes and can mediate transport between vascular bundles (Webb and Gorham 1962; Zhang et al. 2010; Zhang et al. 2012; Sui et al. 2021). Despite these interconnections, we did not observe the *S. ureilytica* pathogen within the fascicular phloem, suggesting that the pathogen either did not reach this tissue or was unable to survive in it.

The unique localization of the CYVD pathogen also has implications for disease transmission. Extensive feeding damage by squash bugs and cucumber beetles, coupled with slow EFP blockage, could facilitate entry of pathogen cells into damaged EFP sieve tubes.

Notably, this could bypass the need for a highly specialized transmission mechanism. Squash bugs may transmit the pathogen via salivarian infection, although it is unknown if the CYVD pathogen is circulative in squash bugs, or via a contaminated stylus. Alternatively, squash bugs and cucumber beetles may both transmit the pathogen by depositing contaminated frass into feeding wounds. Such stecorarian transmission is how cucumber beetles spread the cucurbit xylem pathogen *Erwinia tracheiphila* (Saalau Rojas et al. 2015). If CYVD transmission occurs through relatively nonspecific feeding interactions, as suggested by the pathogen’s tropism for phloem tissues not primarily involved in photoassimilate transport (Zhang et al. 2010; Kanvil et al. 2016; Schnieder et al. 2022), CYVD may pose a greater risk than other phloem pathogens of transmission by novel or diverse insect vectors, particularly as climate change and global trade expand the ranges of insects and plants. Whether broadened vector transmission is contributing to the recent geographic expansion of CYVD (Rodriguez et al., 2023; Mphande et al., 2024) remains an open question.

## MATERIALS AND METHODS

### Bacterial strains and standard growth conditions

*S. ureilytica* CYVD strain Z07 was isolated from zucchini (*Cucurbita pepo*) by Elizabeth Little at the University of Georgia, USA (Besler and Little 2017). This strain has natural resistance to tetracycline (Mphande et al. 2025b). *Pseudomonas syringae* pv. syringae strain B728 (hereafter *P. syringae* B728a) (Loper and Lindow 1987) is a mesophyll phytopathogen and thus not adapted to growth in phloem tissues. *P. syringae* and *S. ureilytica* were cultured in Luria-Bertani (LB) medium at 30°C. The antibiotics kanamycin (Km) (50 μg/ml) and tetracycline (Tc) (20 μg/ml) were added as needed.

### Construction of a green fluorescent protein-expressing *S. ureilytica* CYVD strain

Plasmid pP*nptII:gfp* (Stiner and Halverson 2002) contains a fusion between the constitutive *nptII* promoter and *gfp,* encoding a green fluorescent protein (GFP). This plasmid confers kanamycin resistance and replicates in *S. ureilytica* using the broad-host-range pBBR1 replicon; a similar plasmid with a pVS1 replicon (Miller et al. 2000) appeared unable to replicate in Z07. The pP*nptII:gfp* plasmid, hereafter designated pGFP, was mobilized into Z07 via triparental mating using pRK2073 as a helper (Better and Helinski 1983). Colonies were selected on plates containing Km and Tc and verified for fluorescence using a Blue Light Transilluminator (Discovery Scientific Solutions, Phoenix, AZ, USA). The identity of the colonies was confirmed by PCR using primers specific to CYVD strains, A79F (5’-CCAGGATACATCCCATGATGAC-3’) and A79R (5’-CATATTACCTGATGCTCCTC-3’) (Zhang et al. 2005), and genomic DNA extracted using the DNeasy Blood and Tissue Kit (Qiagen, Valencia, CA, USA), as described previously (Mphande et al. 2025b). Fluorescent colonies that had the expected 338-bp fragment (Zhang et al. 2005) were designated Z07(pGFP).

### Plant cultivation

Pattypan summer squash seeds (*C. pepo* cv. Sunburst) (Seedway, Hall, NY, USA.) were planted in autoclaved potting soil (Sunshine® Mix no.1; Sungrow, Agawam, MA, USA) in 1-gallon autoclaved high-density polyethylene pots (Grower’s Solution, Cookeville, TN, USA). Plants were maintained at 28°C, 60-80% relative humidity, and a 16 h/8 h photocycle using fluorescent bulbs (6400K, 54-Watt, 5000 lumens) (AgroBrite; Hydrofarm, Shoemakersville, PA, USA).

Plants were fertilized at planting with 9.2 g Osmocote Plus slow-release granular fertilizer (NPK: 15-9-12) mixed into the top two inches of soil of each pot.

### Plant inoculation

To generate inocula, CYVD strains were grown for 24 h on LB agar (Z07) or LB agar with Km (Z07(pGFP)) and the resulting colonies were suspended in 10 mM phosphate buffered saline (PBS) (1.7 mM KH_2_PO_4_, 10 mM Na_2_HPO_4_, 2.7 mM KCl, and 136 mM NaCl, pH 7.4) to an optical density at 600 nm (OD_600_) of approximately 1.0 (∼10^9^ colony forming units per ml). Plants were inoculated with a needle and syringe (Mphande et al. 2025a) by injecting 300 μl of either bacterial inoculum or PBS into the stem below the cotyledon. Specifically, the stem was first pierced with the needle at two sites, namely at the crown and just below the cotyledon, without injecting inoculum or PBS, as this provided a pressure release for the subsequent inoculum. Second, with the needle at a 45-degree angle and the hole facing downward, the stem was pierced 3 to 4 times below the pilot piercing site near the cotyledon, introducing inoculum or PBS within the stem; with each piercing, the needle was moved gently in and out while slowly releasing the inoculum and the plant was rotated slightly so that injection sites occurred radially around the stem. Approximately 20–30% of the bacterial culture or PBS was retained in the plant. This inoculation method was previously validated as inducing symptoms similar to those induce via herbivory by squash bugs and cucumber beetles (Mphande et al. 2025a,b).

### Microscopy imaging

Inoculated plants from each treatment across multiple independent experiments were destructively sampled 14 d after inoculation for microscopic imaging. Stems and petioles were harvested using sterile razor blades and immediately placed in plastic zip-top bags at 4°C to reduce tissue desiccation before imaging. For petiole imaging, sections were taken from regions near the petiole base. For stem imaging, sections were taken from regions either above the cotyledon or near the crown, avoiding the inoculation sites. Tissues were hand-sectioned with sterile razor blades and mounted on microscope slides in sterile water or PBS + 3% glycerol immediately prior to imaging.

Widefield fluorescence microscopy was performed using an inverted Leica DMi8 microscope equipped with a Leica K5 camera, Leica LED8 light source, and the following dry objectives: objective HC PL FLUOTAR 10x/0.32, objective HC PL FLUOTAR L 20x/0.40 CORR, and objective HC PL FLUOTAR L 40x/0.60 CORR. GFP signals were captured using a 475 nm) LED8 excitation line (10% power) and DFT51010 quad-band fluorescence filter cube (dichroic mirror: 500; emission: 519/25), along with corresponding bright-field images.

Exposure settings were held constant across multiple imaging sessions. Composite images of large tissue regions were generated from tile scans that were stitched together using the ‘Mosaic Merge’ function of the Leica LASX software (v3.7.6.25997).

Confocal laser scanning microscopy was performed using a Zeiss LSM 700 microscope equipped with the following dry objectives: objective EC Plan-Neofluar 5x/0.16, objective Plan-Apochromat 10x/0.45, and objective Plan-Apochromat 20x/0.08. GFP was excited using a 488 nm diode laser line (18% transmission) and detected using a descanned PMT detector (375 V gain). Transmitted-light images were acquired simultaneously with GFP using a transmission PMT detector (250 V gain). Digital scan zoom was varied depending on the desired field of view.

For postprocessing, raw image files (.lif or .czi) were loaded into FIJI (version 2.16.0/1.54p) using Bio-Formats Importer. Brightness and contrast were adjusted for display purposes and images were cropped to regions of interest where appropriate. Maximum-intensity projections of GFP fluorescence were generated from confocal z-stacks using the Z project function (Maximum Intensity) in FIJI. No deconvolution was applied to any image.

GFP fluorescence lifetime was measured on a Leica Stellaris 5 confocal microscope, which was equipped with an HC PL APO CS2 20×/0.75 IMM objective, a white-light laser tuned to 498 nm, and an HyD S detector, using the TauScan function within the LAS X software (v4.8.2.29567).

### CFDA labeling

5(6)-Carboxyfluorescein diacetate (CFDA) (Sigma-Aldrich, Saint Louis, MO, USA) was dissolved in DMSO at a concentration of 25 mM. This stock was then diluted in distilled water to a final working concentration of 0.5 mM CFDA in 2% DMSO. For plant labeling, 400-grit sandpaper was used to gently abrade four well-separated sites on the adaxial surface of a single mature leaf of a 5-week-old Sunburst plant. Working CFDA solution (500 μL) was applied to each abraded area, and the area was covered in plastic wrap to limit evaporation. Control plants were abraded in an identical manner but treated with 2% DMSO in distilled water. After 3 h incubation under plant growth conditions, the adjoining petiole was hand-sectioned and examined by epifluorescence microscopy (Leica DMi8; excitation: 475; emission: 519/25).

### Growth assay in squash EFP exudate

Vascular exudate was harvested from two, healthy 5-week-old laboratory-grown squash (*C. pepo* cv. Sunburst) plants. Following surface sterilization with 70% ethanol, exudate was harvested sequentially from five petioles and the main stem. Specifically, a petiole was first cut near the leaf with an ethanol-sterilized razor blade and exudate was collected from the cut surface using a micropipette until flow ceased (approx. 3 min). A second cut was then made on the same petiole closer to the main stem and collection continued until exudation again ceased. This procedure was repeated for each of five petioles, from leaves 7 through 3, before exudate was collected from between the 2^nd^ and 1^st^ node of the main stem. Exudates were pooled, stored at -70°C, and thawed and filtered (0.22 μm) before use in growth assays; a total of 160 μL of filtered exudate was collected. This exudate was assumed to originate primarily from the extrafascicular rather than fascicular phloem based on biochemical studies of exudate collected in this manner from *C. pepo* (Zhang et al. 2010; Zhang et al. 2012; Lopez-Cobollo et al. 2016). Although fluids from cut cells, the apoplast, and the xylem may have diluted the EFP exudate, this dilution was likely attenuated by performing the collection under continuous bright light to prevent the buildup of water pressure in the roots (Zhang et al. 2012) and collecting exudates from higher leaves, which have been shown to contain little xylem sap based on exudate pH (Supplemental data of Schnieder et al. 2022). Although the exudate was viscous, as expected due the known ability of EFP exudate to form gels, the cell suspensions appeared uniform throughout the experiment.

Replicate overnight cultures of *S. ureilytica* Z07 and *P. syringae* B728a were grown in LB at 30°C. Bacterial cells were washed and resuspended in 10 mM phosphate buffer (PB) to a concentration of ∼4x10^5^ cells/mL, upon which 10 μL cells were mixed with 40 μL filtered exudate or 40 μL PB (negative control) in 200-μL PCR tubes. The tubes were incubated in an upright position at 30°C with continuous orbital shaking for 30 h. At 0, 10, and 30 h, a 2.5-μL aliquot was removed from each of two replicate cultures per treatment, and serial dilutions were spot-plated in triplicate onto LB agar. Colony forming units were enumerated after 26 h of incubation at 30°C and imaged at 26 and 50 h.

### Growth assay in the presence of cucurbitacin

Cucurbitacin B (AdipoGen Life Sciences, Cat No AG-CN2-0472) and Cucurbitacin E (AdipoGen Life Sciences, Cat No AG-CN2-0474) were dissolved in methanol at a concentration of 1 mg/mL prior to dilution in LB broth or R2A broth (Neogen, MI, USA) to the desired working concentrations. Cells from two replicate overnight cultures of *S. ureilytica* Z07 and *P. syringae* B728a grown in LB at 30°C were used to inoculate a flat-bottom 96-well microplate (Costar, NY, USA) containing LB or R2A broth with or without cucurbitacin (150 μL per well total), with a starting bacterial density of ∼7x10^7^ cells/mL and cucurbitacin concentrations of 10, 1, 0.1, 0.01, 0.001 and 0 μg/mL. The microplate was incubated in a plate reader (Synergy H1; Biotek, VT, USA) at 30°C for 20 h with continuous double orbital shaking, and the OD_600_ values were collected every 60 min.

## DATA STATEMENT

The data supporting the findings of this study are available within the article and its supplementary materials.

## ACKNOWLEDGEMENTS

We thank several people at Iowa State University for their contributions to this project, including Margaret Carter and Damilola Olatunji for assistance with confocal microscopy, Nick Peters for assistance with epifluorescence microscopy, Larry Halverson for providing plasmid pP*nptII:gfp*, and Lynn Clark for assistance with identifying plant anatomical features in microscopy images. This work was funded by the Organic Research and Extension Initiative (OREI) of USDA-NIFA, Grant No. 2019-51300-30248, and by USDA-NIFA Hatch Project No. IOW05814.

## SUPPORTING INFORMATION

**Figure S1.** Annotated schematics indicating the major vascular and nonvascular tissues in (a) stem and (b) petiole sections of *Cucurbita pepo*.

**Figure S2.** Images showing that Z07(pGFP) infection causes disease symptoms comparable to those caused by wild-type Z07 in *C. pepo*.

**Figure S3.** Growth data showing that *Serratia ureilytica* strain Z07 and *Pseudomonas syringae* pv. syringae B728a grow well in *C. pepo* EFP exudate and are not inhibited by cucurbitacins.

**Figure S4.** Microscopy images indicating that Z07(pGFP) may also localize to intercellular spaces adjacent to the EFP.

