## Supporting Information for "The bacterial pathogen causing cucurbit yellow vine disease associates with the specialized extrafascicular phloem unique to cucurbits"

**Figure S1.** Annotated schematics indicating the major vascular and nonvascular tissues in (a) stem and (b) petiole sections of *Cucurbita pepo.*

**Figure S2.** Images showing that Z07(pGFP) infection causes disease symptoms comparable to those caused by wild-type Z07 in *C. pepo*.

**Figure S3.** Growth data showing that *Serratia ureilytica* strain Z07 and *Pseudomonas syringae* pv. syringae B728a grow well in *C. pepo* EFP exudate and are not inhibited by cucurbitacins.

**Figure S4.** Microscopy images indicating that Z07(pGFP) may also localize to intercellular spaces adjacent to the EFP.

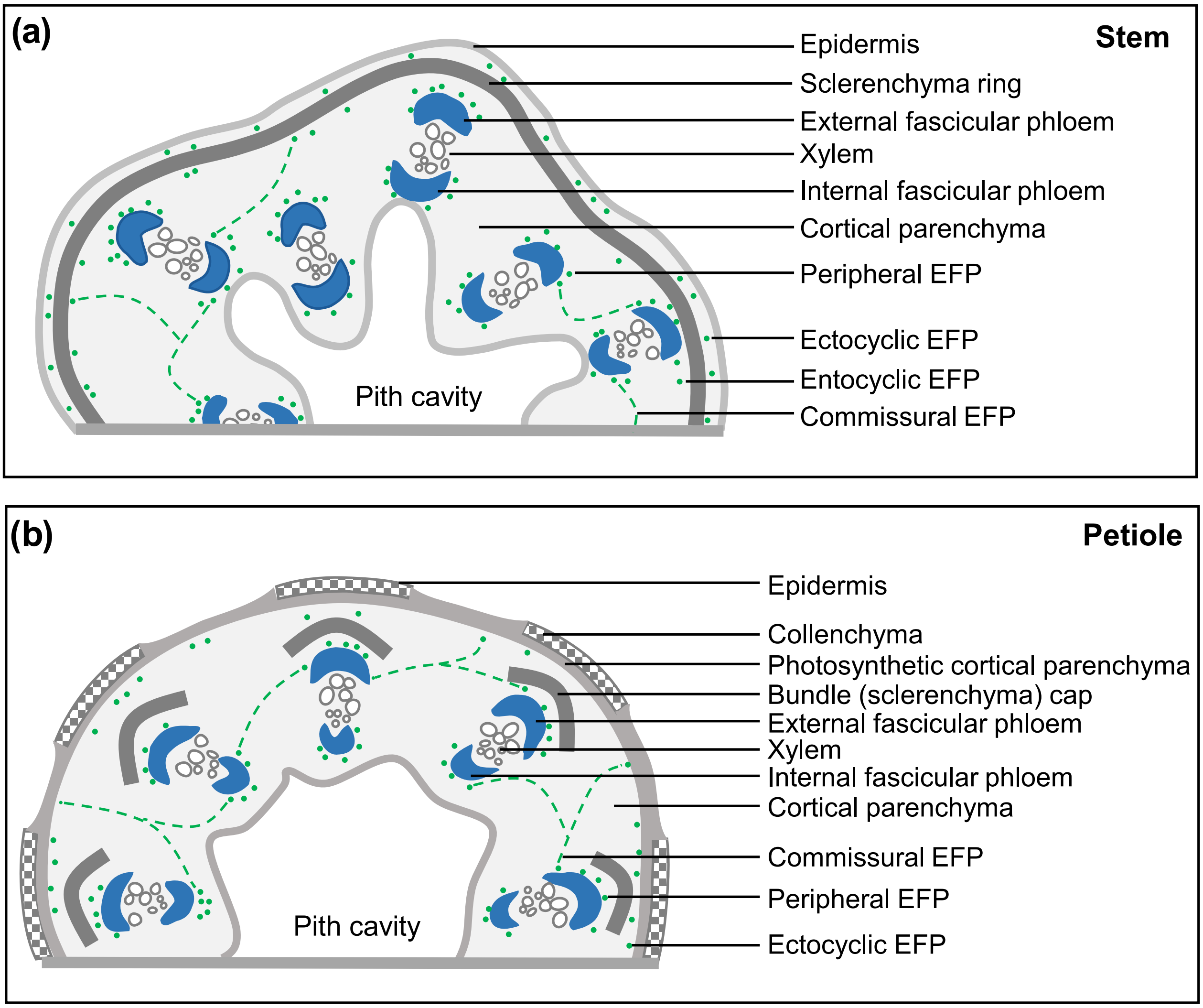


**Figure S1. Annotated schematics indicating the major vascular and nonvascular tissues in transverse (a) stem and (b) petiole sections of *Cucurbita pepo.*** Fascicular (bundle) phloem is colored blue, and extrafascicular phloem (EFP) types are colored green. The ectocyclic and entocyclic EFP cannot be formally differentiated in petioles due to the absence of a sclerenchyma ring. We designate these as ectocyclic EFP, as they are located external to the sclerenchyma fibers that make up individual bundle caps and in regions without sclerenchyma fibers lie immediately interior to the ring of photosynthetic cortical parenchyma.

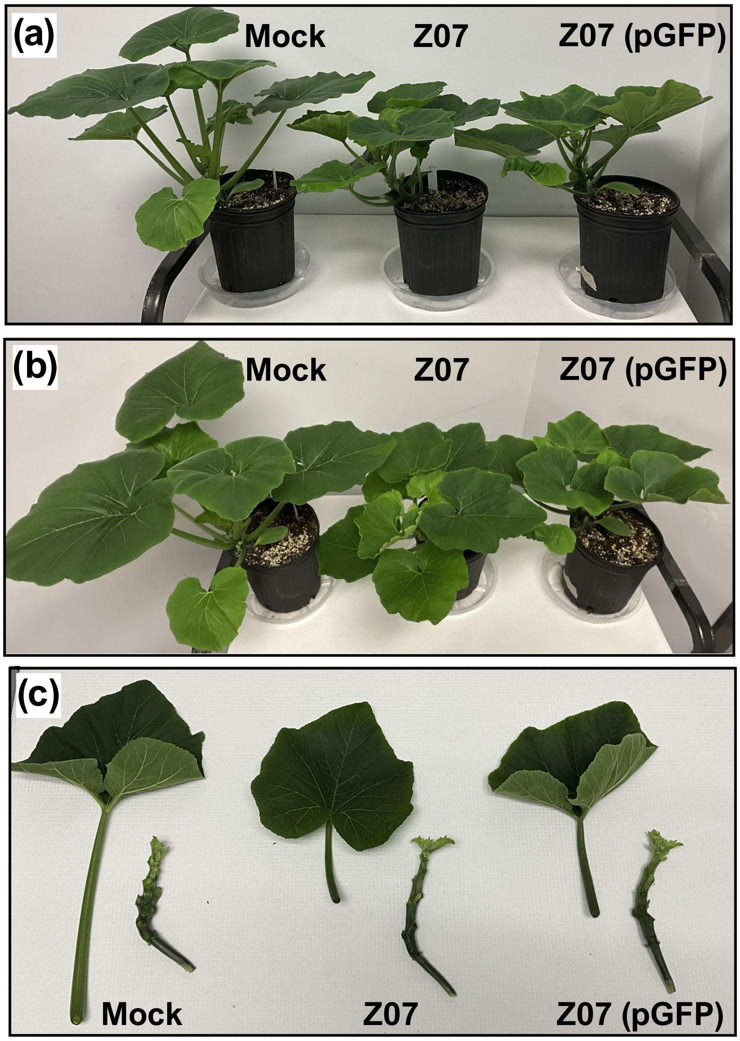


**Figure S2. Z07(pGFP) infection causes disease symptoms comparable to those caused by wild-type Z07 in *C. pepo*.** Pictures show squash cultivar Sunburst at 2 weeks after needle-inoculation with buffer (Mock) or cell suspensions of strain Z07 or Z07(pGFP). (a, b) CYVD-infected plants exhibit leaf cupping and appear stunted compared to the mock-inoculated control plant. (c) CYVD-infected plants have elongated stems (left) and shortened petioles (right) compared to the mock-inoculated control plant, as seen in Mphande et al. (2025a).

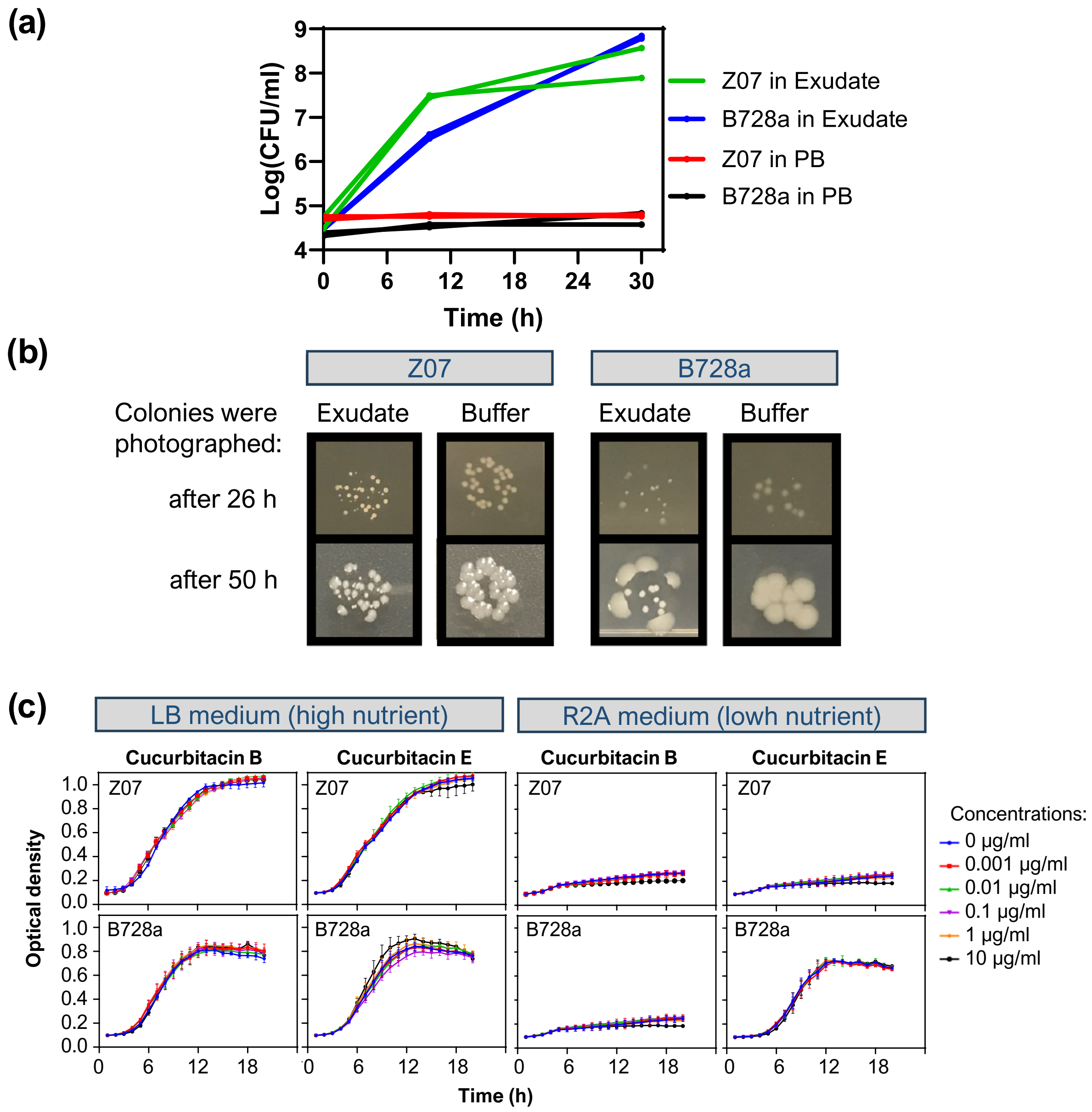


**Figure S3.** ***Serratia ureilytica*** **strain Z07 and** ***Pseudomonas syringae*** **pv. syringae B728a grow well in** ***C. pepo*** **EFP exudate and are not inhibited by cucurbitacins.** (a) Growth of bacteria in squash exudate or phosphate buffer (PB) over time as monitored by plating for colony-forming units (CFU). Data are shown for each of two replicate cultures per treatment. (b) Colony morphology of bacteria following introduction into squash exudate and immediate plating onto an LB plate without exudate; exudate induced heterogeneous colony morphology of Z07 and suppressed apparent swarming motility of B728a. (c) Growth of bacteria in the presence of either cucurbitacin B or E as amended into the high-nutrient medium LB or the low-nutrient medium R2A. Values shown are the mean optical density at 600 nm ± standard deviation ($n = 2$).

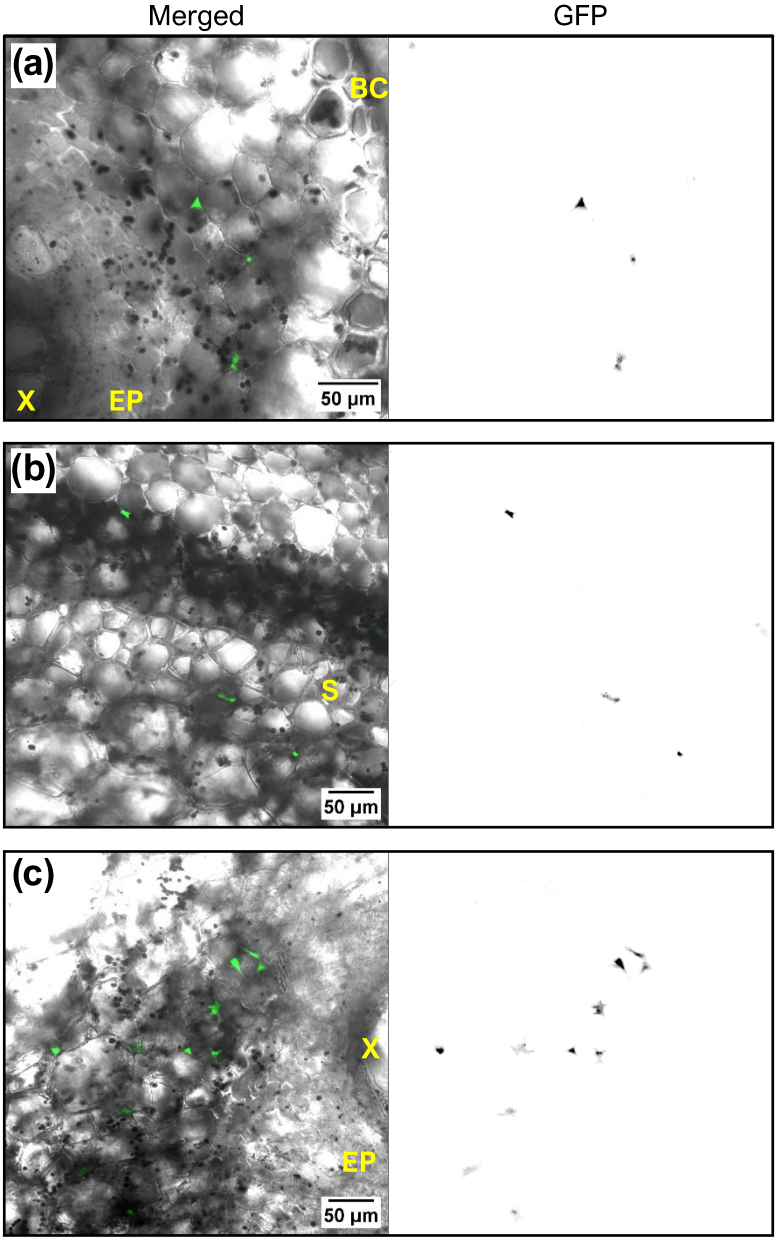


**Figure S4. Z07(pGFP) may also localize to intercellular spaces adjacent to EFP.** Merged transmitted light/GFP confocal microscopy images and corresponding GFP channel alone of *C. pepo* cross-sections. For all panels, the GFP channel alone is shown as an inverted grayscale image. (a) Petiole region spanning from the xylem (X) to the sclerenchyma bundle cap (BC). EP, external fascicular phloem. (b) Stem region encompassing the sclerenchyma ring (S). (c) Stem region encompassing the xylem (X), external fascicular phloem (EP), and adjacent parenchyma cells.
